# The effect of common inversion polymorphisms *In(2L)t and In(3R)Mo* on patterns of transcriptional variation in *Drosophila melanogaster*

**DOI:** 10.1101/128926

**Authors:** Erik Lavington, Andrew D. Kern

## Abstract

Chromosomal inversions are an ubiquitous feature of genetic variation. Theoretical models describe several mechanisms by which inversions can drive adaptation and be maintained as polymorphisms. While inversions have been shown previously to be under selection, or contain genetic variation under selection, the specific phenotypic consequences of inversions leading to their maintenance remain unclear. Here we use genomic sequence and expression data from the Drosophila Genetic Reference Panel to explore the effects of two cosmopolitan inversions, *In(2L)t* and *In(3R)Mo*, on patterns of transcriptional variation. We demonstrate that each inversion has a significant effect on transcript abundance for hundreds of genes across the genome. Inversion affected loci (IAL) appear both within inversions as well as on unlinked chromosomes. Importantly, IAL do not appear to be influenced by the previously reported genome-wide expression correlation structure. We found that five genes involved with sterol uptake, four of which are Niemann-Pick Type 2 orthologs, are upregulated in flies with *In(3R)Mo* but do not have SNPs in LD with the inversion. We speculate that this upregulation is driven by genetic variation in *mod(mdg4)* that is in LD with *In(3R)Mo*. We find that there is little evidence for regional or position effect of inversions on gene expression at the chromosomal level but do find evidence for the distal breakpoint of *In(3R)Mo* interrupting one gene and possibly disassociating the two flanking genes from regulatory elements.

## Introduction

Chromosomal inversions, in which a portion of linear DNA sequence is flipped in its orientation, are a common member of the menagerie of DNA polymorphisms, and have been found in diverse organismal populations such as humans, plants, and fruit flies (KRIMBAS and POWELL 1992; KIDD et al. 2010; LOWRY and WILLIS 2010). In many cases, large chromosomal inversions have profound impacts on phenotype and disease (FEUK 2010). For instance recurrent inversions are responsible for an estimated 43% of hemophilia A cases (LAKICH et al. 1993). Inversions can also have beneficial effects. A 900kb inversion on human chromosome 17 (q21.31) has been shown to be associated with higher female fecundity in the Icelandic population (STEFANSSON et al. 2005). In populations of the malaria vector An. gambiae a large chromosomal inversion on chromosome 2L (2La) is associated with desiccation resistance and thus segregates at high frequencies in arid environments (FOUET et al. 2012). These examples are the very tip of the iceberg—inversion polymorphisms have been implicated in numerous phenotypic differences among a host of organisms, however little is known about the mechanisms by which inversions confer their phenotypic effects.

Perhaps the single best studied inversions are those from Drosophila, in part made famous by the pioneering work of Dobzhansky (DOBZHANSKY and STURTEVANT 1938). Dobzhansky focused much attention on spatial and temporal variation in frequency of large inversions of D. pseudoobscura and showed in broad strokes that clear fitness differences were responsible for the regular patterns of frequency change observed. These findings in turn spurred a large body of population genetics theory to explain the establishment and selective persistence of inversions in natural populations (LEVENE and DOBZHANSKY 1958; FRASER et al. 1966; ANDERSON et al. 1967; TOBARI and KOJIMA 1967). As postulated by Sturtevant (1921), crossover suppression induced in inversion heterozygotes can mean that a single adaptive allele within an inversion may suffice for the selective invasion of that rearrangement (Haldane 1937). Such lowered levels of recombination and attendant increases in linkage disequilibrium (LD) could thus present the opportunity for subsequent coadaptation of multiple genes near inversion breakpoints (STURTEVANT and MATHER 1938; DOBZHANSKY 1947). Conversely locally adapted alleles that predate the rearrangement on the same chromosome might aid the establishment of an inversion simply because of the reduction in recombination rates between such loci (KIRKPATRICK and BARTON 2006). Further, inversions might have direct fitness effects, for instance by deletion or changes in gene expression near the inversion breakpoints (KIRKPATRICK and KERN 2012). At present we have precious little information as to the variants responsible for differential fitness effects associated with inversions.

In *Drosophila melanogaster* paracentric inversions spanning several megabases are common and have been found in populations across the globe (Stalker 1976, 1980; Knibb *et al.* 1981; Sezgin *et al.* 2004; Anderson *et al.* 2005; Umina *et al.* 2005). Much of this segregating inversion polymorphism is associated with latitudinal clines in *D.melanogaster*, an historically tropical species adapting along tropical-to-temperate climatic gradients in Australia and North America (KNIBB *et al.* 1981; WEEKS *et al.* 2002; DE JONG and BOCHDANOVITS 2003; SEZGIN *et al.* 2004; REINHARDT *et al.* 2014; SCHRIDER *et al.* 2016). Clinally varying phenotypes that are associated with inversions include heat resistance, cold tolerance, and body size (Weeks *et al.* 2002; Anderson *et al.* 2003; de Jong and Bochdanovits 2003). Clinal variation of inversion frequency in *D.melanogaster* has been shown via population genetic approaches to be due to selection independent of demography (Reinhardt *et al.* 2014; Kapun *et al.* 2016), though migration has been suggested to generate these patterns along with local adaptation (BERGLAND *et al.* 2016). Indeed, inversions have been observed to have a major effect on several phenotypes that vary between temperate and tropical populations across several *Drosophila* species and these clines have been stable since their discovery roughly 80 years ago (HOFFMANN *et al.* 2004; COGNI *et al.* 2017). Unfortunately, while the associations are known, the molecular mechanisms at work determining differential phenotypes as a result of inversion status are still unknown.

Recent population genomic projects in *D.melanogaster*, such as the Drosophila Genetic Reference Panel (DGRP) and Drosophila Population Genomics Project (DPGP), have opened the opportunity to study inversions systematically as these resources have captured segregating inversions from North America and Africa (POOL *et al.* 2012; MACKAY *et al.* 2012; LANGLEY *et al.* 2012; HOULE and MÁRQUEZ 2015). Corbett-Detig *et al.* (2012) bioinformatically mapped previously unknown breakpoints of several inversions, a task that was tedious for even single inversions prior to whole genome sequencing (WESLEY and EANES 1994; ANDOLFATTO *et al.* 1999; MATZKIN *et al.* 2005). For instance Corbett-Detig *et al.* (2012) discovered the breakpoints associated with numerous inversions and demonstrated the expected increase in LD near inversion breakpoints and elevated differentiation between inverted and standard arrangement chromosomes at the nucleotide level. In parallel with the exponential increase in population genomic resources, large-scale phenotypic association studies of these same genotypes have been accumulating (MACKAY et al. 2012; VONESCH et al. 2016; TELONIS-SCOTT et al. 2016). These include numerous phenotypes previously associated with inversion polymorphism such as body size (WEEKS et al. 2002) and desiccation resistance (HOFFMANN et al. 2005).

A logical place to look for inversion effects that may influence suites of phenotypes would be transcript level variation. Previous findings strongly suggest that inversions could be important drivers of adaptation with gene expression variation as a potential molecular mechanism (CHAMBERS 1991; LÓPEZ-MAURY *et al.* 2008; FRASER 2013). Indeed inversions could affect patterns of transcript variation in a number of ways: **1)** genes at or near inversion breakpoints may become disabled or separated from their regulatory apparatus, thus inversions may have direct effects on transcription, **2)** increased LD in inversions due to crossover suppression may increase linkage with gene expression Quantitative Trait Loci (eQTL), and thus alternative alleles of the inversion may be associated with differential expression of genes within the inversion (i.e. indirect, cis-eQTL associated with the inversion), **3)** eQTL in LD with the inversion might themselves regulate genes outside of the inversion (i.e. indirect, trans-eQTL associated with the inversion), or **4)** the large-scale nature of *Drosophila* inversions may create global changes in the organization of chromatin or nuclear localization of the chromosomes such that genes are differentially regulated between inversion and standard karyotypes. Thus inversions may have a direct effect on global patterns of transcription, and act as trans eQTL themselves. Indeed earlier studies of transcriptional variation in *D.melanogaster* have hinted at the influence of inversions on genome wide patterns of gene expression (AYROLES *et al.* 2009; MASSOURAS *et al.* 2012; HUANG *et al.* 2015). Here we address the effect of two cosmopolitan inversions, *In(2L)t* and *In(3R)Mo*, on patterns of transcription by using whole-genome sequence, gene expression, and inversion call data from the DGRP.

## Methods/Materials

### Materials

Processed expression data previously reported in (Ayroles et al. 2009) was downloaded from ArrayExpress (KOLESNIKOV et al. 2015). We accepted inversion state calls for each of the DGRP lines where cytological and bioinformatic inversion calls for In(2L)t and In(3R)Mo agree and removed lines from analyses where there was any disagreement (CORBETT-DETIG et al. 2012a; HUANG et al. 2014; HOULE and MÁRQUEZ 2015). Using the same databases, we removed Individuals from lines likely heterozygous for In(2L)t or In(3R)Mo. Expression analyses were performed with 34 lines (136 individuals), with two lines homozygous for In(2L)t and seven lines homozygous for In(3R)Mo. There were 26 lines homozygous for the Standard arrangement at both In(2L)t and In(3R)Mo, as one line was homozygous for both inversions. We calculated LD between In(2L)t and SNPs using 181 lines including 19 inversion bearing lines. We calculated LD between In(3R)Mo and SNPs using 197 lines including 17 inversion bearing lines. To maintain consistency with Affymetrix library files, dm3/BDGP release 5 genomic coordinates and annotations corresponding to BDGP version 5.49 were used in conjunction with the Affymetrix Drosophila 2 Release 35 library file update.

### Methods

Statistical analyses were performed in R (R CORE TEAM 2013) using the functions (`lm`), (`anova`), (`quantile`), (`qvalue`), (`phyper`), and (`ggplot2`).

### 3’ UTR Array Analysis

*Correlation structure:* Pairwise gene expression correlation coefficients were calculated by linear regression on all unique pairwise combinations of probe sets, excepting self-comparisons. Correlation coefficients reported here as adjusted r^2^ from the R function (`lm`). Gene expression modules used here were reported in Ayroles et al (2009). Null distributions of uniquely occupied clusters for each inversion were generated by permuting the occupied cluster for each gene 100,000 times and calculating the total unique clusters occupied by inversion affected loci (IAL) for each permutation.

*Inversion effect on expression:* Linear regressions of sex, inversion, and Line effects with expression as the response were performed for each probe set:

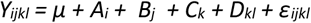

for individual expression value, *Y_ijkl_*, in response to *i*^th^ Sex, *A_i_*, *j*^th^ *In(2L)t* state, *B_j_*, *k*^th^ *In(3R)Mo* state, *C_k_*, and the *l*^th^ Line in the *j*^th^ *In(2L)t* state, *D_kl_*, with *ε_ijkl_* as the error term in Lines. Model testing was performed using (`add1`) and (`drop1`) in R to add interaction terms or remove main effect terms from the above model, respectively. Interaction terms were added one at a time to the main effects and tested for each probe set. An AIC is reported for each model with a lower absolute value being preferred when comparing two models. The effect of adding an interaction term between Sex and *In(2L)t* or Sex and *In(3R)Mo* varied by probe set, but the above model was the best fit for 10,082 of the probe sets. Models with and without an interaction term between inversions performed the same for all loci thus we chose the less complex model. Similarly, the above model was the best fit for 12,994 probe sets when compared to dropping any of the main effect terms. We calculated *p*-values of the observed *F* values as percentiles of the *F* distributions generated by 10,000 permutations sampling each inversion independently without replacement. Multiple testing correction was performed by calculating *q*-values using the R package (`qvalue`) with FDR=0.05 (STOREY and TIBSHIRANI 2003) on the permutation-derived *p*-values. Proportion of variance explained by each effect was calculated as η^2^. Magnitude and direction of inversion effect was calculated as Cohen’s d (COHEN 1988). Cohen’s d is a description of the difference between two standardized distributions and the expected proportion of overlap between the distributions can be estimated given a false positive rate. For example, for two distributions, each with a standard deviation of 1, a Cohen’s d of 1 represents a difference of 1 standard deviation between the means and a ~62% expected overlap given a false positive rate of 5%.

### Functional Annotation Enrichment

Functional annotation profiling was performed using the g:Profiler online portal of g:GOSt using default settings (REIMAND *et al.* 2016) (version r1622_e84_eg31). Ambiguous 3’UTR probe sets were resolved manually if possible, or ignored if they overlapped transcripts for more than one gene.

### SNP-Inversion LD

Linkage disequilibrium (LD) was calculated as r^2^=D’/p_S_(1-p_S_)p_I_(1-p_I_) for each diallelic SNP S and inversion I, with major allele frequencies p_S_ and p_I_, using a custom bash script. For each SNP, significance was calculated as a chi-squared (χ^2^) transformation with 1 degree of freedom of r2 as χ2=Nr2, where N is the sample size. Significant LD was defined as a SNP with sample size of at least 60, minor allele frequency of 10% for both the SNP and inversion state, and χ2 > critical value (p=0.05, d.f.=1) with Bonferroni correction for all SNPs on that chromosome arm (n=967774, χ2 > 29.65329 and 947970, χ2 > 29.61321 for chr2L and chr3R, respectively). The sample size and minor allele frequency cutoffs ensured that there were at least six representative lines bearing minor alleles. Genes were considered in significant LD with inversion state if at least one significant SNP was found within the annotated gene region (FlyBase v5.49).

### IAL physical clustering

To see if inversion affected loci (IAL) were physically clustered within the genome, we examined physical clustering by measuring the coefficient of variance (CV) of distances between genes by chromosome arm. Location and length of each gene was used from FlyBase v5.49. Distance between neighboring genes was calculated as the distance between ends of gene annotated regions of neighboring genes. Distance from the most distal or proximal genes to the distal or proximal endpoint, respectively, was not included. CV was calculated as the standard deviation divided by the mean of the distribution of intergenic distances for each chromosomal arm (SOKAL and ROHLF 1995). We defined the intergenic distance between overlapping gene regions as zero. Null distributions of intergenic distances for each chromosome and each inversion were generated by 100,000 random samples, without replacement, of the same number of genes from a chromosome arm as the number of IAL for that chromosome arm and inversion, then calculating the distances between those genes. Confidence intervals were calculated as 2.5%-97.5% quantiles from the corresponding random sample distribution.

### Transcription factor-target gene interactions

Transcription factors (TFs) and target genes (TGs) were defined by Drosophila Interaction Database modMine (CONTRINO *et al.* 2012) (v2015_12). Genes in this analysis were those that are present in Affymetrix Drosophila2 genome array annotation (Release 35), DroID TF-TG database v2015_12, and FlyBase v5.49 annotation. Over/under-representation of genes with significant inversion effect on expression as targets of transcription factors with SNPs in LD with inversion state was calculated as the probability of the observation given the hypergeometric distribution.

Custom scripts, data, and analysis results can be found online at https://github.com/kern-lab/lavingtonKern, including file descriptions in the AnalysisFiles.readme document.

## Results

To examine what influence, if any, common inversion polymorphisms have on patterns of transcription in the Drosophila genome we combined publically available genome sequences (MACKAY *et al.* 2012), their associated karyotype calls (CORBETT-DETIG *et al.* 2012a; HUANG *et al.* 2014; HOULE and MÁRQUEZ 2015), and previously published microarray based expression data(AYROLES *et al.* 2009). We validated the use of a model with only main effects of Sex, Line, and two cosmopolitan inversions, *In(2L)t* and *In(3R)Mo* using the R functions (add1) and (drop1). This main effects model performed better than any model having any of the main effect terms removed or with the addition of any interaction term (see Methods). After correction for multiple testing (see Methods), we found 229 and 498 total probe sets with significant inversion effects for *In(2L)t* and *In(3R)Mo*, respectively, hereafter referred to as inversion affected loci (IAL) (Figure 1). These IAL occur both within the inversions themselves (40 *In(2L)t*, 111 *In(3R)Mo*), outside the inversions but near the breakpoints (3 *In(2L)t*, 38 *In(3R)Mo* within 1Mb of the breakpoint), as well as scattered throughout the genome (134 *In(2L)t*, 181 *In(3R)Mo*; see Table 1). This is a large number of loci with transcript abundance variation correlating with inversion state; however, we note that the inversion effect contribution to variance is relatively small for the vast majority of loci (Supplemental data).

**Figure 1.**
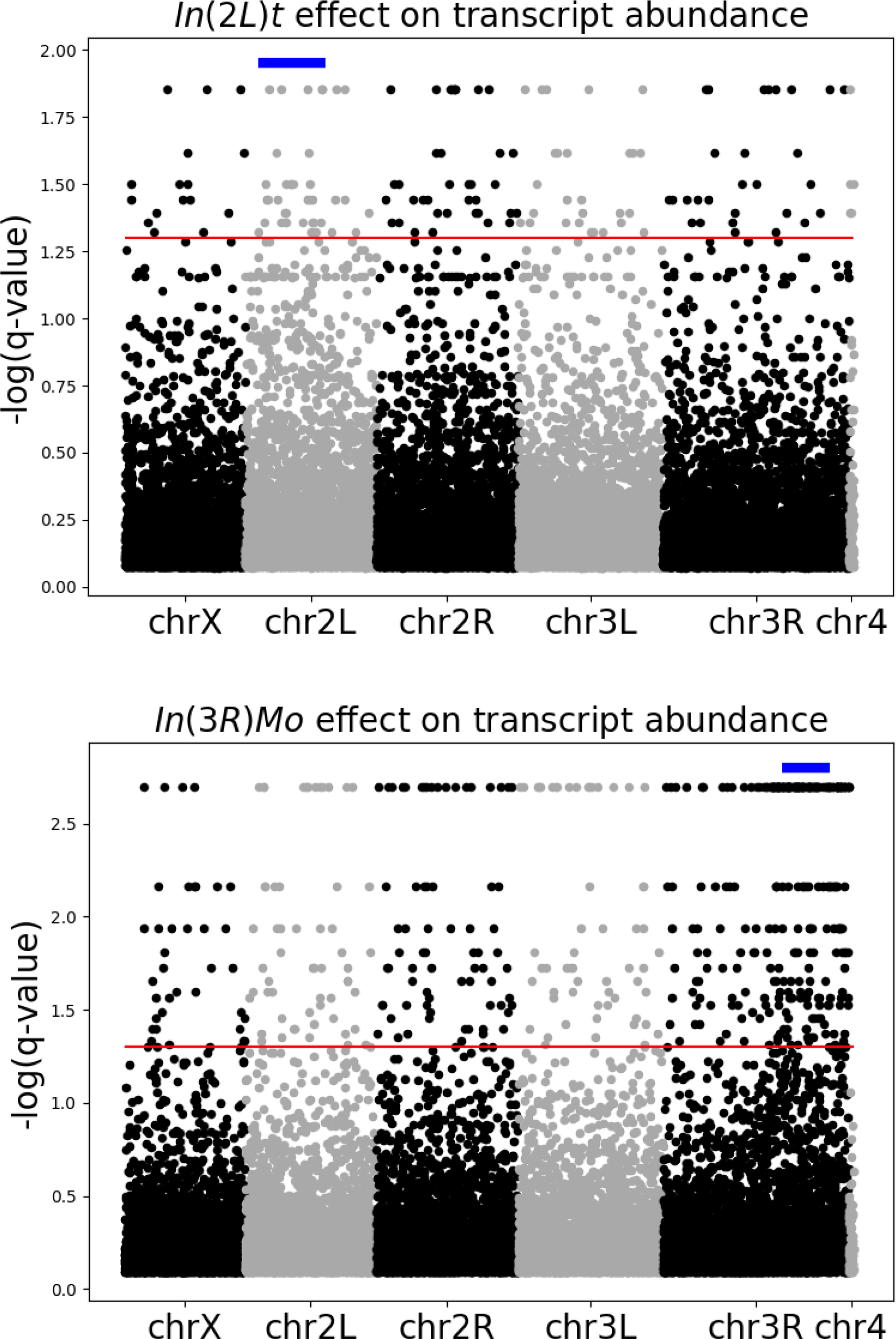
Manhattan plot of log transformed, Bonferroni corrected p-values for an *In(2L)t* (top) and *In(3R)Mo* (bottom) inversion effect on transcription for each probe set across the genome. The blue bar above the points indicates the genomic location of the inversion. The red horizontal line represents the genome-wide significance threshold of p=5x10-8

**Table 1.** Location of IAL. Counts of IAL between breakpoints, on the same chromosome as the inversion but outside of the breakpoints, and those on the same chromosome as the inversion but outside, and within 1Mb, of the breakpoints, as well as total IAL on each chromosome arm for each inversion.

|  | Within<br>Inversion | Outside<br>Inversion, on<br>same Chr | < 1Mb<br>outside<br>Inversion | Chr 2L | Chr 2R | Chr 3L | Chr 3R | Chr X | Chr 4 |
| --- | --- | --- | --- | --- | --- | --- | --- | --- | --- |
| ***In(2L)t*** | 40 | 18 | 3 | 61 | 42 | 36 | 35 | 17 | 4 |
| ***In(3R)Mo*** | 111 | 133 | 38 | 44 | 58 | 49 | 244 | 30 | 0 |

One explanation for the large number of IAL found across the genome is that few loci are directly affected by the inversion and the remaining loci are affected indirectly by an expression variation correlation structure, previously described by Ayroles et al (2009). We addressed this correlation structure by the numbers of unique expression modules occupied by, and the distribution of correlation coefficients of, IAL as compared to all genes. If a significant portion of the IAL we observe are due to expression variation correlation, then we would expect that IAL occupy fewer expression modules than the same number of genes drawn at random. We would also expect the mean correlation of IAL between IAL to be higher than the genome wide average. We observe IAL occupy more modules than expected at random for both *In(2L)t* (71 obs; 38-56 95% c.i.) and *In(3R)Mo* (108 obs; 65-87 95% c.i.). For both inversions, we did not observe higher mean correlation between IAL between IAL and non-IAL or the genome-wide mean (Table 2).

**Table 2.** Pairwise correlation of gene expression. Mean r^2^ was calculated from all pairwise correlation for the set description. IAL x IAL is the set of all pairwise correlation coefficients of expression between IAL for that inversion. IAL x other is the set of correlation coefficients of expression between IAL and all other (non-IAL).

| | | Mean $r^2$ |
| --- | --- | --- |
| *In(2L)t* | IAL x IAL | 0.0965 |
|  | IAL x other | 0.0962 |
| *In(3R)Mo* | IAL x IAL | 0.1025 |
|  | IAL x other | 0.1042 |
|  | **Genome-wide** | 0.1181 |

We then examined the inversion effect on gene expression variation for four distinct categories of effect: 1.) *cis-* or 2.) *trans*-inversion effects of SNPs in LD with the inversion, 3.) direct effects of the inversion by interrupting genes, and 4.) regional effects of chromosomal rearrangement.

## *Cis*-Inversion effect of SNPs in LD with the inversion

Our model explicitly tests the effect of chromosomal arrangement on expression variation and here we focus on SNP variation as the main driver of the inversion effect by taking advantage of LD between the inversion state and SNP variants. LD with inversions is highest at the breakpoints and decays in both directions from each breakpoint as expected (Figure 2) (WESLEY and EANES 1994; ANDOLFATTO *et al.* 2001; LANGLEY *et al.* 2012; CORBETT-DETIG *et al.* 2012b). To determine *cis*-effects of the inversions we tested for over-representation of IAL among loci in LD with inversion state. This ignores the location of the loci and focuses on the correlation of SNP alleles with the inversion state. As expected, few loci in LD with the inversion were located on a different chromosomal arm (3 with *In(2L)t*, and 2 with *In(3R)Mo*)) and IAL in LD are located only on the same chromosomal arm (Table 3). We observed a significant overrepresentation of IAL with SNPs in LD with inversion state (Table 4).

**Figure 2.**
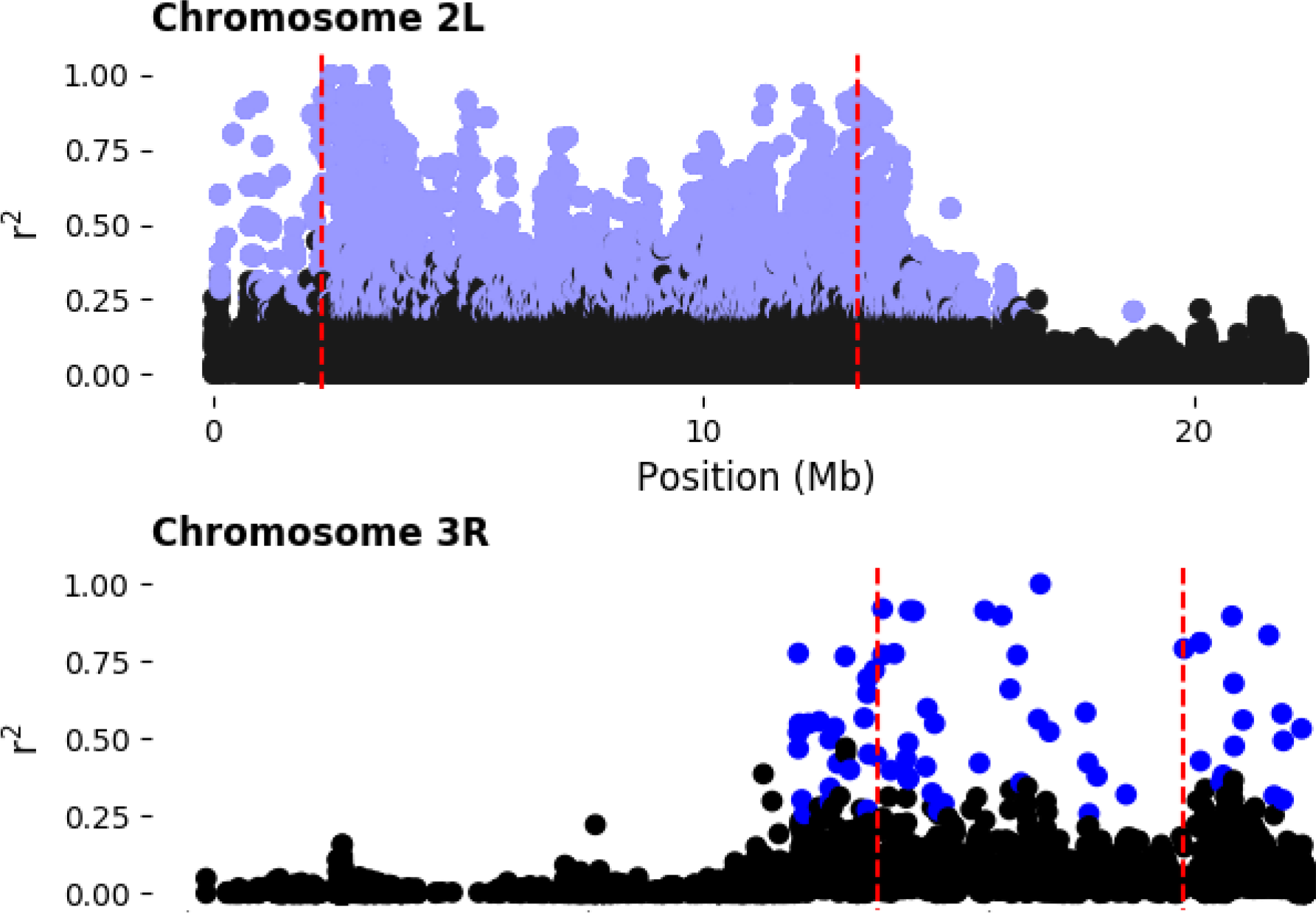
Linkage disequilibrium between SNPs and inversion state. Disequilibrium is measured as r^2^. Inversion breakpoints are depicted as red vertical lines. SNPs with at least 10% minor allele frequency, for Inversion and SNP, and are in significant LD with the inversion are shown in light blue (p<0.05, Bonferroni correction for all SNPs detected by chromosome arm).

**Table 3.** Location of loci with SNPs in LD with inversions. Loci are identified as BDGP v5.49 and Affymetrix library v.35 annotated gene regions containing SNPs in significant LD with the inversion (see Methods).

|  |  | chr2L | chr2R | chr3L | chr3R | chrX |
| --- | --- | --- | --- | --- | --- | --- |
| *In(2L)t* | All loci | 370 | 1 | 0 | 1 | 1 |
|  | IAL | 11 | 0 | 0 | 0 | 0 |
| *In(3R)Mo* | All loci | 0 | 2 | 0 | 663 | 0 |
|  | IAL | 0 | 0 | 0 | 110 | 0 |

**Table 4.** IAL in LD with inversion. Expected counts (E[X]) and p-values are calculated by hypergeometric distribution with uncorrected p-value cutoff of p=0.025 for the two-tailed test. Genes included in analyses are annotated in both BDGP v5.49 and Affymetrix library version 35.

| **Inversion** | Total Genes | Total IAL | IAL in LD with Inversion | Genes in LD with Inversion | E[X] | p(E[X]>x) |
| --- | --- | --- | --- | --- | --- | --- |
| ***In(2L)t*** | 11969 | 192 | 11 | 372 | 6 | 0.01683 |
| ***In(3R)Mo*** | | 425 | 102 | 744 | 26 | $6.08 \times 10^{-35}$ |

## *Trans*-inversion effect of SNPs in LD with the inversion

Our expectation of a *cis*-inversion effect is dependent on SNP variation in LD with the inversion. By the same rationale, a *trans*-inversion effect may be detected as an IAL without SNP variation in LD with the inversion, as we observe with a majority of IAL for each inversion (181 of 192 for *In(2L)t* and 323 of 425 for *In(3R)Mo* (see Table 4). Assuming SNP variation is the basis of expression variation, one trivial explanation of a *trans*-inversion effect is SNP variation in transcription factors in LD with the inversion acting on downstream targets. The TFs in this case need not be IAL as SNPs in protein coding regions of TFs can give rise to expression variation in downstream targets. However, we found no over- or underrepresentation of IAL that are targets of TFs with SNPs in LD with either inversion (Table 5). A possible example of *trans*-inversion effect was found by functional analysis of IAL and discussed below.

**Table 5.** IAL as targets of Transcription Factors with SNPs in LD with inversion. Expected counts E[X] and p-values calculated by hypergeometric distribution with uncorrected p-value cutoff of p=0.025 for the two-tailed test. Genes included in these analyses are annotated in BDGP v5.49, Affymetrix library v.35, and DroIDb v2015_12.

| **Inversion** | Total Genes | DroID TFs with Sig SNPs | Total DroID TF targets | Total DroID TF targets also IAL | Targets of Sig SNP DroID TFs also IAL | E[X] | p(E[X]>x) |
| --- | --- | --- | --- | --- | --- | --- | --- |
| ***In(2L)t*** | 10623 | 3 | 3802 | 181 | 53 | 65 | 0.96260 |
| ***In(3R)Mo*** |  | 8 | 2373 | 385 | 99 | 86 | 0.04801 |

## Direct effect of inversions by disrupting genes at breakpoints

Nucleotide sequence variation at each breakpoint of an inversion can truncate transcribed gene regions and disassociate transcribed regions from transcription factor binding sites and other regulatory elements. Truncation most likely leads to down-regulation of genes and rearrangement of regulatory elements and can lead to up- or downregulation of genes at either breakpoint. The closest IAL to any of the four breakpoints are near the distal breakpoint of *In(3R)Mo*. *CG1951* and *βGalNAcTB* are within 2.5kb inside and outside of the inversion region, respectively, and *Ssl2* is interrupted by the breakpoint. All three are downregulated in the presence of the inversion. The next closest IAL is 24 kb away. The closest IAL to the proximal breakpoint of *In(3R)Mo* are 32kb away. It is also important to note that loci immediately surrounding the proximal breakpoint are not transcriptionally affected by the rearrangement, so the presumed disassociated regulatory elements from the distal end are not altering expression of loci at the proximal end. The closest IAL to the proximal and distal breakpoints of *In(2L)t* are 34kb and 37kb away, respectively, and thus probably too far to have been affected by direct effects of the breakpoint.

## Regional effect of chromosomal rearrangement

We tested for regional effects of chromosomal rearrangement by looking for over- or underrepresentation of genes in LD with the inversion as IAL. We did observe more IAL than expected in LD with each inversion (Table 4) and a trend of transcriptional downregulation across the *In(2L)t* region (Figure 3 A & D), but note that the effect size is relatively small. Near the breakpoints, patterns of inversion effect direction are generally small and not likely significantly different from zero (Figure 3 B,C,E & F). The pattern of downregulation immediately surrounding the distal breakpoint of *In(3R)Mo* is moderate and appears to be driven by three loci: *CG1951, Ssl2*,and *β4GalNAcTB*. This is likely a direct effect of the inversion on *Ssl2*, as the breakpoint interrupts this gene proximal to *CG1951* (CORBETT-DETIG *et al.* 2012a), and possibly disassociation of regulatory elements from coding sequence with respect to *CG1951* and *β4GalNAcTB*.

**Figure 3.**
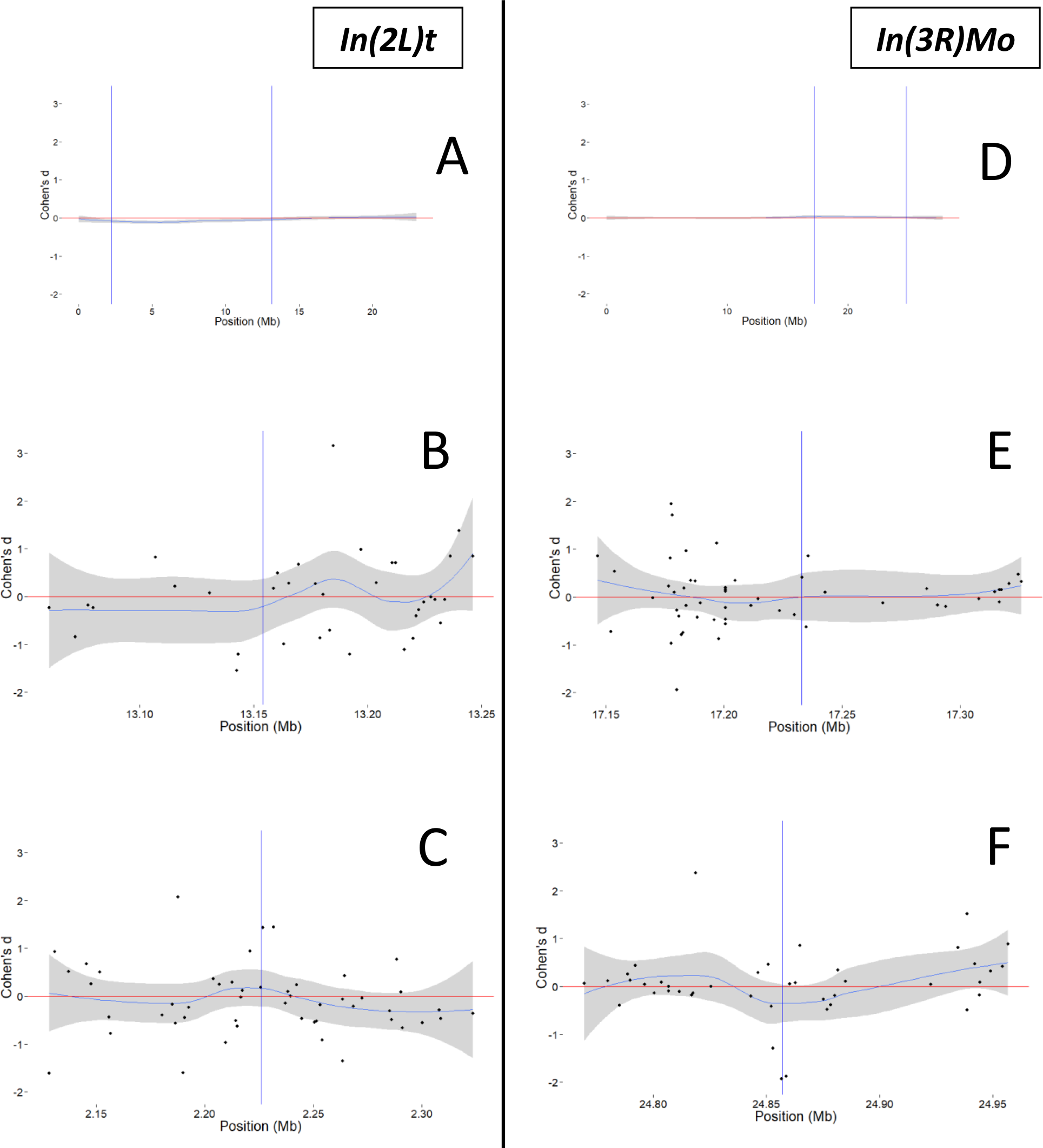
Cohen’s d of inversion effect for probe sets by location. Cohen’s d for *In(2L)t* effect across chromosome 2L (A) and for *In(3R)Mo* effect across chromosome 3R (D). Loess curves (polynomial line) and 95% confidence intervals (shaded area) for 200kb surrounding each breakpoint: *In(2L)t* (B) proximal and (C) distal, *In(3R)Mo* proximal (E) and distal (F). Inversion breakpoints are depicted as vertical lines. Loess curves and confidence intervals generated by (geom_smooth) function in R package (ggplot2). Positive values of Cohen’s d represent increased transcript levels associated with the inverted chromosome state and negative values represent decreased transcript levels.

We also examined whether IAL tend to cluster together by physical location along chromosomes, which could arise from more localized regional effects. We measured physical clustering as the coefficient of variation (CV) of distances between genes, for the global CV, and between IAL by chromosome arm. For each arm, IAL for both inversions were less clustered than the distribution of all genes, although *In(3R)Mo* IAL are more clustered on chromosome 3R than we would expect for the same number of randomly drawn genes, but are still less clustered than the genomic background generally (Table 6).

**Table 6.**
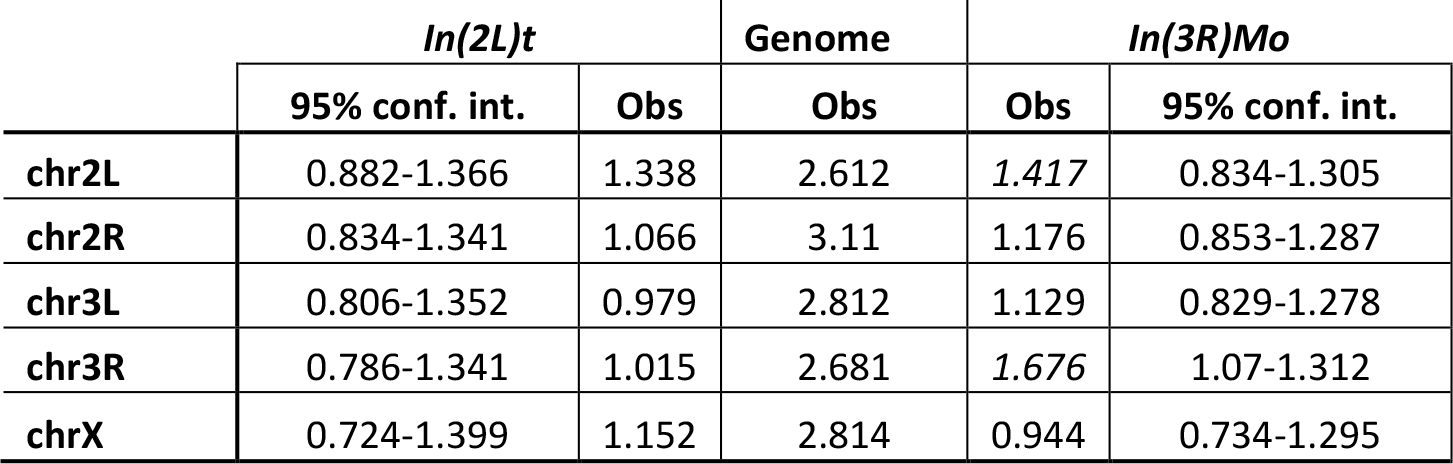
Gene location dispersion as described by the Coefficient of Variation (CV). Observed genome CV’s calculated from loci in both Flybase ver.5.49 and Affymetrix Drosophila2 r35 annotation databases by chromosome arm. See Methods for details.

## Functional analysis

Coadapted alleles segregating with an inversion should also be in LD with the inversion breakpoints. We used gProfiler g:GOSt functional profiling to detect overrepresentation of functional groups in sets of IAL. The sets of IAL that we analyzed were those IAL in LD with the inversion (Table 2) or targets of another gene with variation segregating with the inversion (Table 4). Functional analysis of IAL for each inversion yielded significant groups only when considering all *In(3R)Mo* IAL or only IAL where inversion state explains at least 15% of expression variance (Supplemental data). Sterol transport is significant in both cases (*p*=0.022 for all, *p*=0.000116 for ≥15% variance) and catalytic activity term (GO:0003824) is significant when considering all IAL(p=0.000146). We found no significant functional groups when considering any similar grouping of *In(2L)t* IAL.

The sterol transport group found to be enriched among *In(3R)Mo* IAL includes four of the eight Niemann-Pick type 2 orthologs (*Npc2b*, *Npc2c*, *Npc2f*, *Npc2g*) (HUANG *et al.* 2007), along with *Apoltp*. All five genes are upregulated in association with the inverted arrangement, suggesting an increase of sterol uptake in *In(3R)Mo* bearing flies. In the context of the expression samples, most of these genes are preferentially expressed in the adult gut (modENCODE, Contrino *et al.* 2012). While the four NPC2s are on chromosome 3R (Figure 4), *Apoltp* is on chromosome 2L, and none of these genes contain a SNP in the gene region that is in significant LD with *In(3R)Mo*, or each other, and only *Npc2f* is within the inversion region (Figure 5). It is possible that the significant upregulation of the sterol transport group is a downstream effect of *mod(mdg4)*, which is an IAL near the proximal breakpoint (Figure 4) and in LD with *In(3R)Mo* (Figure 5). *mod(mdg4)* is a chromatin protein associated with *Npc2b* as well as other chromatin proteins and TFs associated with all eight *Npc2* orthologs (Supplemental data). We address this exciting finding in further detail in the Discussion.

**Figure 4.**
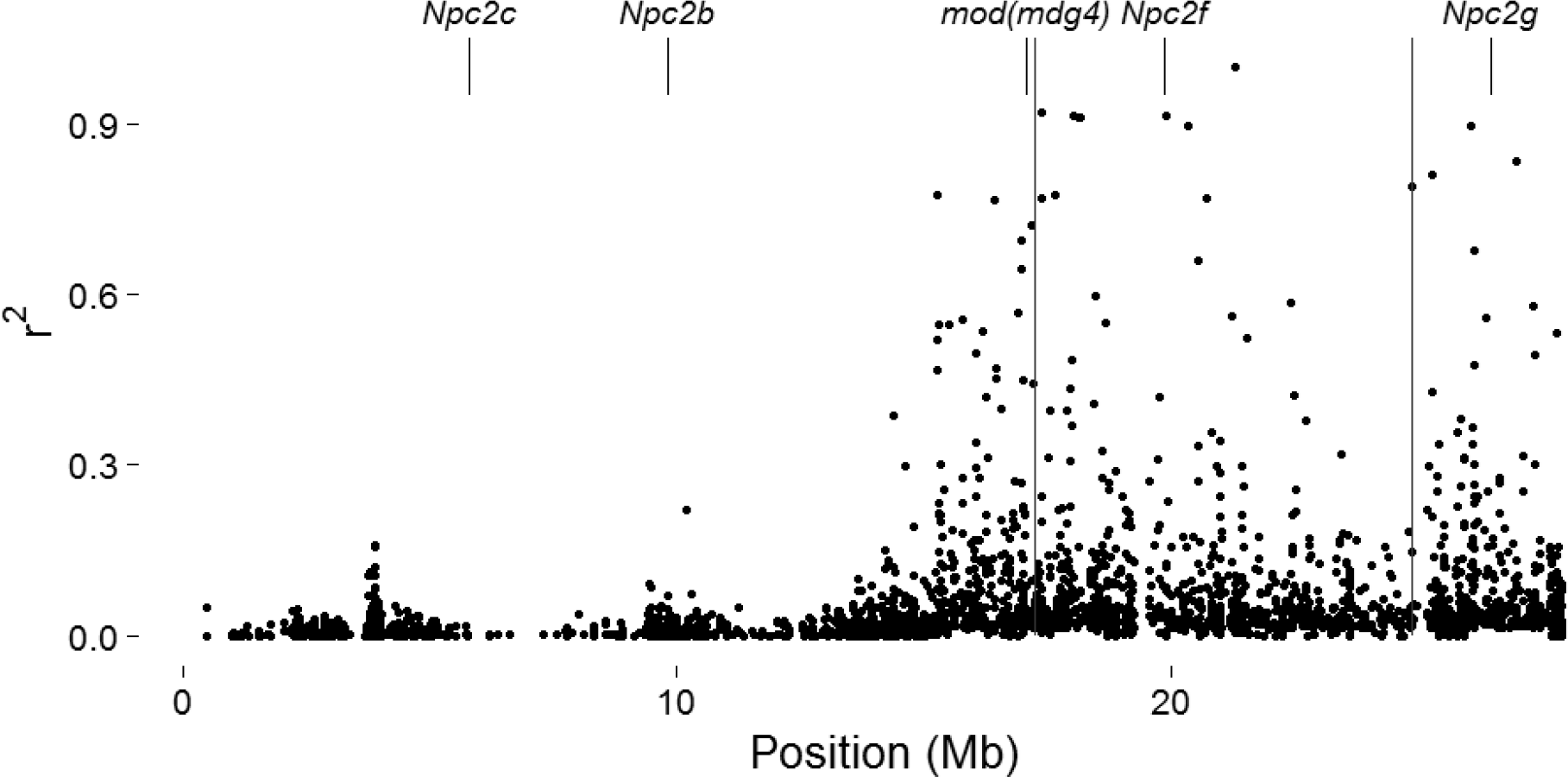
Location of *Npc2s* and *mod(mdg4)* on chromosome 3R. Long vertical lines represent breakpoints of *In(3R)Mo*. Only *mod(mdg4)* contains a SNP in LD with *In(3R)Mo*.

**Figure 5.**
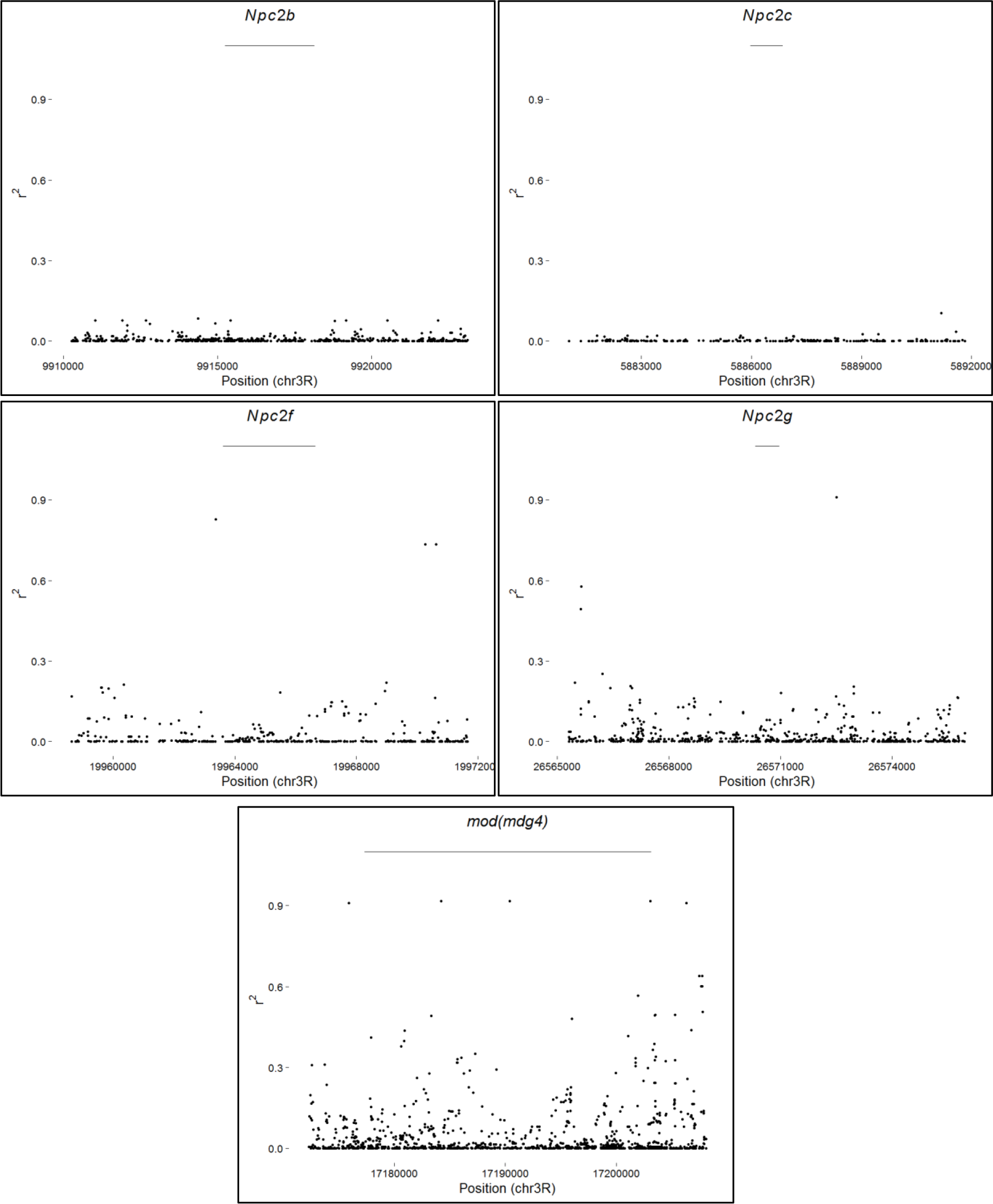
Correlation of SNPs with *In(3R)Mo* by gene. All SNPs within gene regions and within 5kb up- and down-stream of each gene: *Npc2b, Npc2c, Npc2f, Npc2g*, and *mod(mdg4)*. Thin horizontal bars above each plot represent the gene region for that gene.

## Discussion

Despite decades of research on polymorphic inversions in *Drosophila melanogaster*, and despite an overwhelming consensus that inversions are maintained due to selection, we have little understanding of the targets of selection in these inversions that leads to their maintenance (cf. Kirkpatrick and Kern 2012). Here we examine the role of transcriptional variation induced by cosmopolitan inversions to explore what effect, if any, inversions might have on gene expression that itself might be selectively favored. Besides gross rearrangement effects, we examined gene disruption at inversion breakpoints, IAL in LD with the inversion, and IAL not in LD with the inversion. We assumed that multiple functionally linked IAL to be potentially coadapted so long as they contained SNPs in LD with the inverted arrangement. We also assumed that *trans*-inversion effects, IAL not in LD with the inversion, could be the result of these loci interacting with TFs in LD with the inversion or epistatic interactions with loci in LD. We note that while sample size of *In(2L)t* bearing individuals in the expression analysis is small (8 of 136), we were still able to detect a relatively large number of significant loci across the genome. We believe that our methods were conservative and limit the number of false positives at the expense of a likely high number of false negatives. We argue that we are accounting for the small sample sizes in our analyses and interpretations. We also note that we could not address over- or under-dominance in this study as we examined only lines known to be homozygous for either arrangement of *In(2L)t* or *In(3R)Mo*.

Of particular concern was what role, if any, expression correlation had on our findings. To explore this we considered pairwise correlation coefficients and previously described expression modules (AYROLES *et al.* 2009). One could imagine that the true false discovery rate would be much higher than anticipated due to expression correlation of multiple loci from the same expression module. We found that IAL for both inversions occupied more expression modules than we would expect when drawing genes at random. Even when considering just the pairwise expression correlation, we found that IAL for both inversions were slightly less correlated on average than the genome wide mean. These observations suggest that the inversion effects arose independent of the underlying correlation structure.

We found many loci of significant effect, yet most have a modest contribution to expression variation. Importantly, the only evidence of direct structural influence on patterns of expression was at a single breakpoint that appears to be due to the inversion mutation interrupting gene regions but not moving genes to a different region of the chromosome. That is, there was no appreciable pattern of up- or down-regulation of genes along the chromosome with respect to the location of inversion breakpoints. Perhaps unsurprisingly, we did find an overabundance of expression perturbation within the inversion regions themselves. This would suggest the effect is possibly due to genetic, rather than the structural, variation in LD with the inversion. That is, transcription variation associated with the inversion is due to genetic variation at the gene level and not the rearrangement of loci.

Inversion polymorphism can be maintained by reduced recombination between locally coadapted alleles within inversions (DOBZHANSKY and OTHERS 1970), both with (NEI *et al.* 1967; PEPPER 2003) or without epistasis (KIRKPATRICK and BARTON 2006). Clinal inversion variation in *D.melanogaster* within *In(3R)Payne* in Australian populations and *In(3R)Payne, In(2R)NS, and In(2L)t* in American populations has been shown to be due to selection rather than demographic history and indicative of coadaptation (KENNINGTON *et al.* 2006; KOLACZKOWSKI *et al.* 2011; KAPUN *et al.* 2016). We reasoned that one might be able to detect epistatic coadaptation through functional analysis of IAL. Coadaptation of non-interacting loci would likely result in either no significant functional groups or multiple, unrelated significant groups. We found functional annotation enrichment only when we considered all IALs with respect to *In(3R)Mo*. This would suggest that either one locus or a few coadapted, but non-interacting, loci segregate with these inversions to maintain polymorphic inversions. However, it is still possible that coadapted loci could be overlooked in our analysis; certainly the existing annotation is incomplete. Moreover, our statistical power to detect IAL suffers due to the constraints of the number of inversions captured in the DGRP dataset. Nevertheless our observation that there are multiple IAL within the inversion with argues strongly for the role that inversions play as modifiers of recombination which may hold adaptive haplotypes together.

Our strongest functional finding, that sterol uptake associated with *In(3R)Mo*, appears to be driven by genetic variation in a single locus as a *trans-*inversion effect. Four of the five genes in this cluster are located on chromosome 3R, however none of these genes have a SNP in significant LD with *In(3R)Mo*, or with each other (Supplemental data), and only one is found within the inversion region (*Npc2f)*. This would rule out effective coadaptation of these genes and suggest that the location of these genes on the same chromosome as the inversion is coincidence. Assuming the upstream effector of the sterol transport is a transcription factor, it could contain a SNP that alters either its protein-coding sequence or expression. Either scenario would require that the genetic variant responsible for the upregulation of the four Npc2s in question would have to be in LD with *In(3R)Mo*. Only one TF, *mod(mdg4)*, annotated in DroIDb as interacting with any (*Npc2b*) of the five sterol transport IAL contains a SNP in LD with *In(3R)Mo*. Furthermore, *mod(mdg4)* is itself an IAL and also located near the proximal breakpoint of *In(3R)Mo*. We speculate that inversion associated expression variation detected in this functional group is under control of *mod(mdg4)*.

Increased sterol uptake fits nicely with the positive correlation of *In(3R)Mo* frequency with latitude (KAPUN *et al.* 2014). *Npc2* genes control sterol homeostasis via uptake of dietary sterols in *D.melanogaster*(HUANG *et al.* 2007; CARVALHO *et al.* 2010; NIWA *et al.* 2011). Two species of *Drosophila* differentially express *Npc2’s* in response to cold acclimation, though the patterns differ between the two (PARKER *et al.* 2015). Increased dietary cholesterol increases cold tolerance in *Drosophila melanogaster* (SHREVE *et al.* 2007), however, *D.melanogaster* takes up phytosterols more efficiently than cholesterol (COOKE and SANG 1970). Furthermore, *D.melanogaster* is a sterol auxotroph (CARVALHO *et al.* 2010; NIWA *et al.* 2011) that utilizes dietary sterols preferentially to biosynthesizing different sterols from dietary sterols (CARVALHO *et al.* 2012). This would suggest that *In(3R)Mo* carrying *D.melanogaster* could be cold-acclimated due to increased uptake of dietary sterols, rather than the upregulation of cholesterol production.

It is difficult to interpret results for *In(2L)t* without a clear functional annotation group associated with the inverted state. Assuming *In(2L)t* is under selection (KAPUN *et al.* 2016), the simplest interpretation is that *In(2L)t* polymorphism is maintained by only a small number of loci in LD with the inversion. It is possible that one or more loci in LD with *In(2L)t* contain protein coding variation under selection and no appreciable transcript abundance variation with respect to chromosomal arrangement. We note that there are only 14 IAL in LD with *In(2L)t* (Supplemental data). While a lack of a significant functional group may be dissatisfying, this does provide a manageable candidate list for validation of single targets.

## Conclusion

We found that two different cosmopolitan inversions in *D.melanogaster* have some effect on the expression of hundreds of genes across the genome. While we caution that our sample sizes, particularly for *In(2L)t*, are very small, the permutation approach that we have taken is conservative. The genetic variation responsible for the observed transcriptional variation is only in small part due to the inversion event itself, with the majority of the variation being the result of either allelic variation in LD with the inversions, trans-effects of regulators that also are in LD with the inversions, or as of yet uncharacterized, indirect effects of the inversions. Our results mirror those of a recent report on transcriptional variation caused by inversions in *Drosophila pseudoobscura* (FULLER *et al.* 2016). Fuller et al. demonstrated quite convincingly that the well-studied polymorphic inversions of *D. pseudoobscura* are modulating levels of transcription at multiple life history stages and even found hints of trans-effects. However, due to experimental design constraints they could not examine inversion effects on unlinked chromosomes. Our findings extend this pattern to *Drosophila melanogaster* and show that inversions are affecting loci genome-wide. While we have begun to parse the possible causes of IAL, our focus on available data limits the statistical power of our study. Thus it will be important to conduct carefully designed experiments on inversion polymorphism in *D.melanogaster* to elucidate the true extent of influence of inversions on genome-wide patterns of transcription.

## Acknowledgements

We thank Dan Schrider and Steve Schaeffer and two anonymous reviewers for feedback on the manuscript. A.D.K. and E.L. were supported in part by NIH award no. R01GM078204.

## Supplemental Methods

**Pairwise LD between SNPs**: was calculated with a custom Linux script as r^2^ and χ^2^ calculated as χ^2^ = Nr^2^, as described in Methods. We used the DGRP and excluded lines where either *In(2L)t* or *In(3R)Mo* was suspected to be segregating, We included SNPs that fell within the specified gene regions as annotated in FlyBase v5.49. SNPs with no base calls across all tested lines were excluded. Significance level was corrected for 843,051 tests (all unique, LD calculations for 1299 SNPs).

**Supplemental Table 1.**
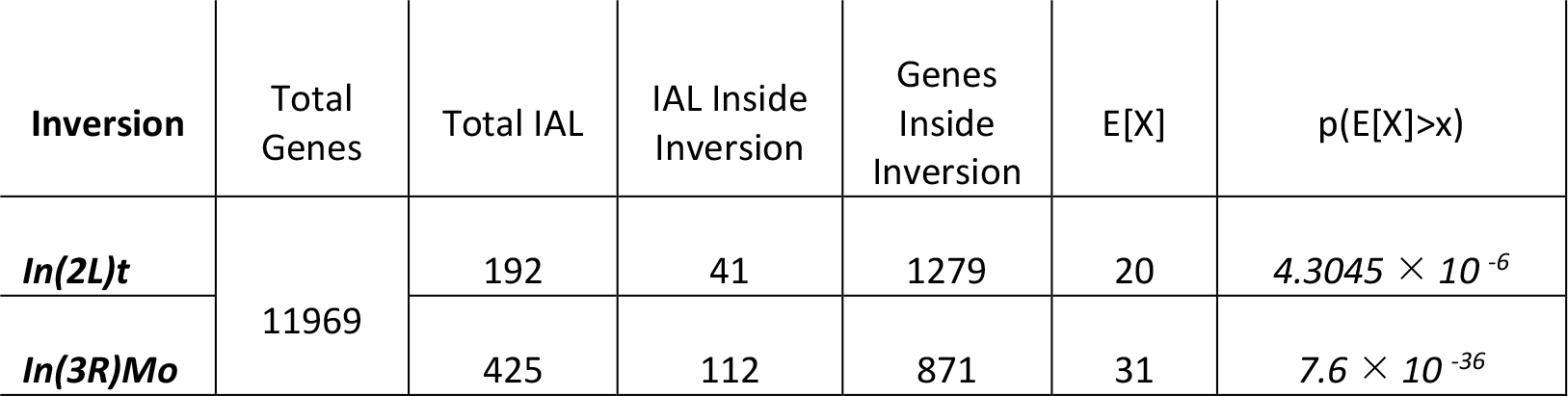
IAL inside inversion region. Expected counts E[X] and p-values are calculated by hypergeometric distribution with uncorrected p-value cutoff of p=0.025 for the two-tailed test. Genes included in analyses are annotated in both BDGP v5.49 and Affymetrix library version 35.

**Supplemental Table 2.**
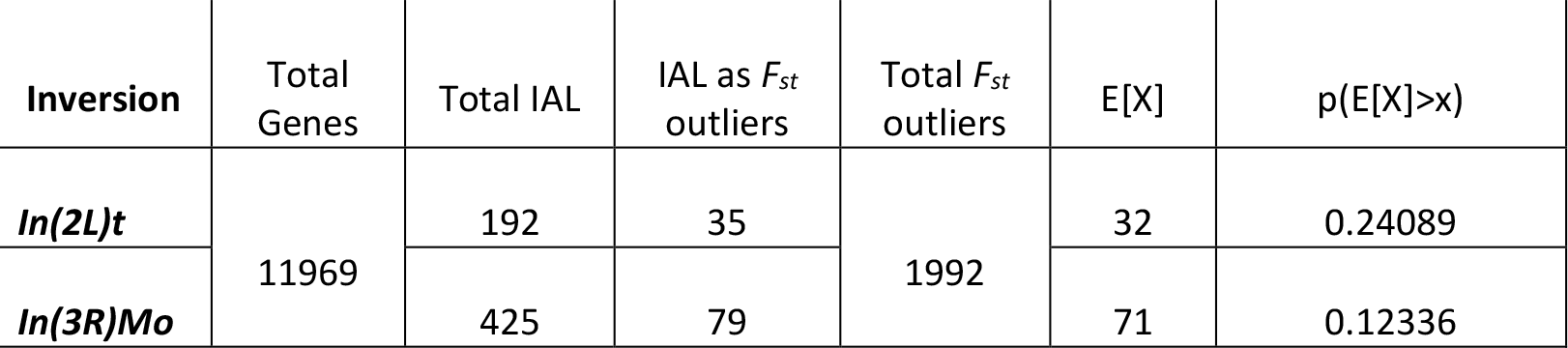
IAL as *F_st_* outliers in clinal populations. Populations from Reinhardt et al (2014). Number of genes reflect the overlap between the unique Affymetrix Drosophila 2 annotated genes and FlyBase r5.49 annotations. Outliers are from the p=0.05 tail of the empirical distribution of 1kb windows (see Reinhardt et al (2014)).

| Inversion | Total Genes | Total IAL | IAL as $F_{st}$ outliers | Total $F_{st}$ outliers | E[X] | p(E[X]>x) |
| --- | --- | --- | --- | --- | --- | --- |
| *In(2L)t* | 11969 | 192 | 35 | 1992 | 32 | 0.24089 |
| *In(3R)Mo* |  | 425 | 79 |  | 71 | 0.12336 |

